## Supplementary Information for "DNA-MGC+: A versatile codec for reliable and resource-efficient data storage on synthetic DNA"

#### Supplementary Notes

##### Supplementary Note S1: Detailed marker-based offset estimation for inner MGC+ code

Supplementary Note S1 extends the offset estimation framework originally introduced in Ref.[1], where the MGC+ code (the inner code of the DNA-MGC+ codec) was developed and analyzed in the *binary* domain. While the overall marker-based synchronization and trellis-based decoding principles remain conceptually the same, the analysis below reformulates the system in the *DNA* domain. The key differences arise from (i) base-level insertion, deletion, and substitution (IDS) errors in the underlying channel model, which modify the transition probability structure, and (ii) the explicit derivation of received DNA marker transition probabilities under quaternary alphabet statistics (Table A).

**Drift and Trellis Formulation** For marker-based encoding, a fixed DNA marker (AC) is inserted periodically after every $\ell_{\text{in}}$ DNA symbols. At the decoder, synchronization is recovered by constructing a trellis whose states represent the cumulative synchronization drift after each marked block. Let the trellis state space be defined as $\Omega = \{-\Phi, -\Phi + 1, \ldots, \Phi\}$, where $\Phi$ is an upper bound on the cumulative drift induced by insertion and deletion events. Let $\mathbf{z} = (z_0, z_1, \ldots, z_v)$ denote the drift sequence, where $z_i \in \Omega$ is the cumulative offset after the $i$-th block and $z_0 = 0$. The drift evolves according to $z_i = z_{i-1} + d_i$, where $d_i$ is the offset increment associated with block $i$. The received DNA sequence $\mathbf{y}$ is segmented into blocks according to the hypothesized drift trajectory as

$$\tilde{\mathbf{b}}^{(i)} \triangleq \mathbf{y}_{[(i-1)\ell'+z_{i-1}+1,\, i\ell'+z_i]}, \quad i \in [v], \tag{S1}$$

where $\ell' = \ell + 2$. The offset increment is equivalently expressed as $d_i = z_i - z_{i-1}$. The last $\mu = 2$ nucleotides of each block correspond to the received marker sequence, denoted by $\tilde{\mathbf{m}}^{(i)}$.

**Marker-centric MAP Drift Estimation** Assuming independent and identically distributed IDS errors across nucleotide positions, the joint probability of observing the received sequence and a particular drift trajectory factorizes as

$$\Pr(\tilde{\mathbf{y}}, \mathbf{z}) = \prod_{i=1}^{v} \Pr(\tilde{\mathbf{b}}^{(i)}, z_i \mid z_{i-1}), \tag{S2}$$

with $z_0 = 0$. To reduce decoding complexity, drift estimation is performed using only the marker observations. This leads to the following marker-centric maximum a posteriori (MAP) objective:

$$\hat{\mathbf{z}} = \arg\max_{\mathbf{z} \in \mathcal{Z}} \prod_{i=1}^{v} \Pr(\tilde{\mathbf{m}}^{(i)}, z_i \mid z_{i-1}), \tag{S3}$$

where $\mathcal{Z}$ denotes the set of all admissible trellis paths constrained to the state space $\Omega$. A graphical illustration of the resulting trellis and its admissible transitions is provided in Fig. A.

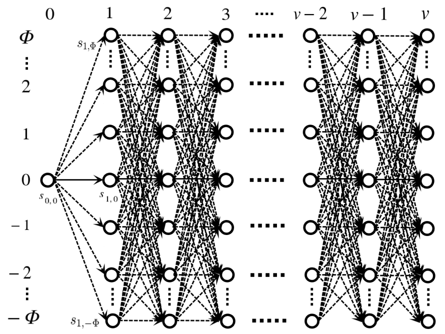

**Figure A.** Trellis representation of the drift sequence $\mathbf{z} = (z_0, z_1, \ldots, z_v)$

**Transition Probabilities** A trellis transition from state $z_{i-1}$ to $z_i = z_{i-1} + d$ is weighted by the probability of observing a received DNA marker $\tilde{\mathbf{m}}$ together with a net offset $d$ over the corresponding block. Since each block consists of a data segment followed by a fixed DNA marker, this probability decomposes as

$$\Pr(\tilde{\mathbf{m}}, d) = \sum_{t=-\mu}^{\mu} \Pr(\mathrm{D}_{\mathrm{data}} = d - t)\ \Pr(\tilde{\mathbf{m}}, \mathrm{D}_{\mathrm{mar}} = t), \tag{S4}$$

where $\mathrm{D}_{\mathrm{data}}$ and $\mathrm{D}_{\mathrm{mar}}$ are random variables representing the cumulative offsets introduced by insertion and deletion events in the data portion and the marker portion of the block, respectively.

The data portion of each block consists of $\ell_{\mathrm{in}}$ DNA nucleotides. Under the standard base-level IDS channel with deletion probability $P_d$, insertion probability $P_i$, substitution probability $P_s$, and $P_r = 1 - P_d - P_i - P_s$, the offset distribution $\Pr(\mathrm{D}_{\mathrm{data}} = t)$ depends only on the block length and channel parameters. Mathematically, it is given by

$$\Pr(\mathrm{D}_{\mathrm{data}} = t) = \sum_{j=\max\{0,-t\}}^{\left\lfloor \frac{\ell_{\mathrm{in}}-t}{2} \right\rfloor} \binom{\ell_{\mathrm{in}}}{j,\ j+t,\ \ell_{\mathrm{in}} - 2j - t} \times P_d^{j}\, P_i^{j+t}\, P_r^{\ell_{\mathrm{in}}-2j-t}. \tag{S5}$$

The marker contribution $\Pr(\tilde{\mathbf{m}}, \mathrm{D}_{\mathrm{mar}} = t)$ depends explicitly on the known transmitted DNA marker $\mathbf{m} = \mathsf{AC}$, the channel parameters, and the received marker sequence $\tilde{\mathbf{m}} \in \{\mathsf{A}, \mathsf{C}, \mathsf{G}, \mathsf{T}\}^*$. Since the marker length is short ($\mu = 2$), this probability can be evaluated exactly by enumerating all possible insertion, deletion, and substitution events affecting the marker nucleotides. For completeness, Table A reports $\Pr(\tilde{\mathbf{m}}, \mathrm{D}_{\mathrm{mar}} = t)$ for all possible received marker sequences $\tilde{\mathbf{m}}$ and offsets $|t| \leq 2$, expressed in closed form as functions of $P_d$, $P_i$, $P_s$, $P_r$ and $P_0$ where $P_0 = 0.25$.

**Dynamic Programming Solution** The MAP problem in (S3) is solved efficiently using dynamic programming. Let $\alpha_i(z_i)$ denote the maximum path metric ending at drift state $z_i$ after processing $i$ blocks. The recursion is

$$\alpha_i(z_i) = \max_{z_{i-1} \in \Omega} \alpha_{i-1}(z_{i-1})\ \Pr(\tilde{\mathbf{m}}^{(i)}, z_i \mid z_{i-1}), \tag{S6}$$

with initialization $\alpha_0(0) = 1$ and $\alpha_0(z) = 0$ for $z \neq 0$. The maximizing predecessor state is stored as

$$\mathrm{pred}_i(z_i) = \arg\max_{z_{i-1} \in \Omega} \alpha_{i-1}(z_{i-1})\ \Pr(\tilde{\mathbf{m}}^{(i)}, z_i \mid z_{i-1}). \tag{S7}$$

After processing all $v$ blocks, the estimated drift sequence $\hat{\mathbf{z}}$ is recovered by traceback starting from

$$\hat{z}_v = \arg\max_{z_v \in \Omega} \alpha_v(z_v), \tag{S8}$$

**Table A.** Marker transition probabilities $\mathbf{Pr}(\tilde{\boldsymbol{m}},\, \mathbf{D_{mar}} = \boldsymbol{t})$ for the DNA marker AC.

| $\tilde{m}$ | $t$ | $\mathbf{Pr}(\tilde{\boldsymbol{m}},\, \mathbf{D_{mar}} = \boldsymbol{t})$ |
|---|---|---|
| AC | 0 | $P_0 P_i P_d + P_r^2$ |
| | +1 | $P_r P_i + (1 - P_d - P_i) P_i P_0$ |
| | +2 | $P_i^2 P_0$ |
| | −1 | $P_0 P_r P_d + P_0^2 P_d P_s$ |
| | −2 | $P_0^2 P_d^2$ |
| AT, AG | 0 | $P_s P_0 P_r$ |
| | +1 | $P_i P_0 P_s$ |
| | −1 | $2 P_d P_0^2 P_s$ |
| | −2 | $P_d^2 P_0^2$ |
| AA | 0 | $P_r P_s P_0 + P_i P_d P_0$ |
| | +1 | $P_i P_s P_0$ |
| | −1 | $P_d P_0 P_r + P_0^2 P_d P_s$ |
| | −2 | $P_d^2 P_0^2$ |
| CC, TC, GC | 0 | $P_0 P_s P_r + P_d P_i P_0$ |
| | +1 | $(1 - P_d - P_i) P_i P_0$ |
| | +2 | $P_i^2 P_0$ |
| | −1 | $P_0 P_r P_d + P_0^2 P_d P_s$ |
| | −2 | $P_0^2 P_d^2$ |
| CT, CG, TT GT, GG, TG | 0 | $P_s^2 P_0^2$ |
| | −1 | $2 P_0^2 P_s P_d$ |
| | −2 | $P_0^2 P_d^2$ |
| CA, TA, GA | 0 | $P_0 P_i P_d + P_s^2 P_0^2$ |
| | −1 | $P_0 P_r P_d + P_0^2 P_s P_d$ |
| | −2 | $P_0^2 P_d^2$ |

and the corresponding offset pattern is obtained as $\hat{d}_i = \hat{z}_i - \hat{z}_{i-1}$, which is subsequently used for inner decoding.

##### Supplementary Note S2: Additional details and comments about some of the comparison codecs

**LDPC Codec** The LDPC-based scheme[2] uses a conceptually different design from the inner–outer architectures adopted by most existing DNA storage codecs. The entire data stream is encoded using a single long binary LDPC code, whose encoded bits are then segmented into short fragments and mapped to DNA symbols. Each fragment carries a BCH-protected index and a marker to assist in handling insertions and deletions during decoding.

In our *in silico* evaluations, we used the implementation provided at `https://github.com/shubhamchandak94/LDPC_DNA_Storage`, keeping its default parameters and setting the `oligo_length` parameter to 150. To obtain different code rates, we replaced the underlying parity-check matrices using three matrices from `https://github.com/shubhamchandak94/LDPC_DNA_storage_data/tree/master/matrices`, resulting in three codec configurations with code rates of 0.58, 0.67, and 0.80 bits/nt. Among these, the configuration with rate 0.58 achieved the best overall performance under the synthetic channel model, as shown in the subsequent Supplementary Tables. However, none of the three configurations was able to decode successfully under the conditions used in the second *in silico* setting.

**DNA-Stairloop** The DNA-StairLoop codec[3] employs a concatenated coding structure consisting of a row code and a column code, but differs from the classical inner–outer architecture through the use of a staircase interleaver that couples the two component codes across adjacent blocks. Decoding follows an iterative soft-information exchange between the row and column decoders. The decoder is designed to operate directly on unclustered sequencing reads and to exploit the presence of multiple reads within its own probabilistic decoding algorithm, without relying on consensus calling. To remain consistent with the decoding procedure proposed by the authors, clustering and alignment were therefore not performed for this codec in our evaluations.

In our *in silico* experiments, we used the reference implementation available at `https://github.com/Guanjinqu/StairLoop`. The parameter `msg_length` was fixed to 19 to obtain encoded DNA sequences of length 147, close to our desired target length of 150. Two values of the parameter `block_num`, 34 and 68, were used to vary the total number of encoded sequences, resulting in two codec configurations with code rates of 0.98 and 0.49 bits/nt, respectively. Our results indicate that this codec is highly sensitive to sequence dropouts under the considered parameters, leading to degraded performance under strong bias conditions. The configuration with rate 0.98 bits/nt outperformed the configuration with rate 0.49 bits/nt, suggesting that increasing the number of encoded sequences while keeping the sequence length fixed does not necessarily improve the performance of this codec. This observation is also implicit in Ref.[3], where the authors evaluate their codec using sequence lengths closer to 200, whereas comparison codecs are evaluated at lengths closer to 150, suggesting that DNA-Stairloop may benefit from allocating more redundancy within individual sequences, and hence operating at longer sequence lengths.

**DNA-Aeon** As reflected in the results in Supplementary Tables S8–S11, the decoding time of DNA-Aeon is significantly higher than that of the other codecs evaluated in this work. Decoding also becomes substantially slower at lower code rates and higher error rates. When the medium-rate configuration of DNA-Aeon was tested at an error rate of 15%, decoding ran for more than two hours without producing a result and was therefore terminated; consequently, no results are reported for this setting. The same occurred for the low-rate configuration at error rates of 10% and 15%.

##### Supplementary Note S3: Discussion about the effect of $\Delta G$ on the coverage distribution

As noted in the Discussion section of the main text, the filtered variants of DNA-MGC+ provide modest but consistent gains in terms of sequencing depth (and read cost) compared to their unfiltered counterparts. This is reflected in the coverage distributions shown in Supplementary Fig. S8, where the unfiltered variants consistently exhibit a substantially larger fraction of underrepresented reference sequences than the filtered variants. For example, under Illumina sequencing for Design B of DNA-MGC+, approximately 200 reference sequences in the unfiltered variant receive between 0 and 10% of the mean coverage, whereas this number decreases to roughly 50 sequences in the filtered variant. The same qualitative behavior is observed across both designs and under both sequencing platforms, demonstrating that filtering systematically reduces the number of poorly covered sequences. These strongly underrepresented sequences contribute to increased sequence dropouts at low sequencing depths, which explains the observed difference in the minimum sequencing depth required for reliable decoding between the filtered and unfiltered variants.

Our analysis of the per-sequence coverage shows that the decoding performance gains of the filtered variants are primarily attributable to the filtering criterion based on the Gibbs free energy $\Delta G$. We support this claim through three complementary statistical analyses of the per-sequence coverage distribution obtained for the unfiltered DNA-MGC+ variants. The numerical results provided below correspond to Nanopore sequencing; qualitatively similar trends were observed under Illumina sequencing.

**Characteristics of underrepresented sequences.** We first identify the subset of reference sequences whose read count falls below 10% of the mean pool coverage. Across the complete unfiltered pool, the mean ± standard deviation (SD) of the maximum homopolymer length, GC content, and $\Delta G$ are $4.2 \pm 1.1$, $48.7\% \pm 3.3\%$, and $-4.1 \pm 2.3$ kcal/mol, respectively. Within the underrepresented subset, the corresponding means are 3.9, 49.9%, and −6.8 kcal/mol, yielding standardized differences from the pool-wide means of −0.27 SD, +0.36 SD, and −1.17 SD. The mean $\Delta G$ of the underrepresented sequences is thus shifted by approximately 1.2 pool-wide standard deviations toward more negative values, which is significant on its own and also substantially larger than the shifts observed for GC content or homopolymer length. Notably, the homopolymer-length shift is small and *negative*, whereas a larger and *positive* shift would be expected if long homopolymers were a major contributor to lower coverage.

**Marginal Spearman correlations.** Spearman correlation analysis further supports the dominant role of $\Delta G$. Across the full unfiltered pool, $\Delta G$ shows the strongest marginal association with per-sequence read count ($\rho = +0.315$, $p$-value $< 10^{-50}$), compared with GC content ($\rho = -0.205$) and maximum homopolymer length ($\rho = +0.082$). The positive sign of the $\Delta G$–coverage correlation confirms that sequences with $\Delta G$ closer to zero (less negative) tend to receive more reads. The weak and positive marginal correlation between maximum homopolymer length and coverage is inconsistent with the hypothesis that long homopolymers cause sequence underrepresentation.

**Partial Spearman correlations.** To verify that the association between $\Delta G$ and coverage is not merely a consequence of its relationship with GC content or homopolymer length, we compute partial Spearman correlations that isolate the $\Delta G$–coverage association after mathematically removing the contribution explained by each of the other variables. The partial correlation between $\Delta G$ and coverage remains largely unchanged when controlling for GC content ($\rho = +0.231$) or for homopolymer length ($\rho = +0.295$), indicating that the association of $\Delta G$ with coverage is largely independent of these two variables. By contrast, the marginal association between GC content and coverage ($\rho = -0.205$) decreases markedly after controlling for $\Delta G$ ($\rho = -0.062$), suggesting that a substantial portion of the apparent GC–coverage association can be attributed to the correlation between GC content and $\Delta G$ rather than to an independent effect of GC content on coverage.

**Summary.** Taken together, these three statistical results indicate that $\Delta G$ is the sequence-level characteristic most strongly associated with coverage variability among those examined here, while GC content and homopolymer length show little to no independent association with sequence underrepresentation in our experiments.

Taken together, these three statistical results indicate that $\Delta G$ is the dominant sequence-level feature associated with coverage variability among those examined here, while GC content and homopolymer length show little to no independent association with sequence underrepresentation in our experiments. A plausible mechanistic explanation is that sequences with more negative $\Delta G$ tend to form stable secondary structures, which may interfere with PCR amplification efficiency and thereby reduce their representation in the sequenced pool. This is also consistent with the observation that the fraction of underrepresented sequences is more pronounced under Illumina sequencing, which involves additional PCR steps during library preparation and cluster generation compared with Nanopore sequencing (see Supplementary Fig. S8).

##### Supplementary Note S4: Comparison with the Gungnir codec

We provide here a comparison between DNA-MGC+ and the recently introduced Gungnir codec[4] under the synthetic error and bias models considered in the main text. Gungnir provides strong IDS error-correction capability by using a complex decoding procedure, in which candidate reconstructions are exhaustively tested until a hash signature associated with the encoded fragment is satisfied. This mechanism gives Gungnir high error tolerance; however, the codec also has two important limitations.

First, as explicitly acknowledged by the authors in Ref.[4], Gungnir does not provide robustness to sequence dropouts.As a result, successful file retrieval requires that each encoded sequence be read at least once. This limitation is architectural and independent of the inner-code IDS error-correction capability, and can therefore substantially affect the required coverage depth and read cost even under ideal inner decoding. Second, Gungnir relies on an exhaustive hypothesis-search procedure whose computational cost becomes substantial at high error rates. For example, the authors report that decoding a 19.4-KB file required more than 37 hours at high error rate on a dual 32-core Xeon Platinum 8369C server. Therefore, to keep the comparison computationally tractable, we evaluate Gungnir on a small, randomly generated 5-KB file at a total error rate of 5%, for the same bias regimes (no bias, moderate bias, strong bias) considered in Fig. 1 of the main text. Unlike the experiments reported in Fig. 1, which restrict each codec to a single CPU core, the Gungnir decoder is allowed to use the full 64-core CPU of our workstation, equipped with an AMD Ryzen Threadripper PRO 7985WX processor. The same computing resources are provided to DNA-MGC+ for a fair comparison.

For reference, we also attempted the no-bias case at a 10% error rate for a 5-KB file, for which Gungnir decoding ran for more than 8 hours without converging while using the full 64-core CPU, exceeding the 2-hour decoding limit adopted throughout our evaluation. In comparison, for the exact same scenario (5-KB file, 10% error rate, no bias), DNA-MGC+ incurs a decoding time of 0.95s at a coverage depth of 16×, and 1.26s at the minimum required coverage depth of 3.75×.

Following the evaluation framework of Fig. 1, we consider three code-rate classes. For Gungnir, the high- and medium-rate configurations are obtained using the Gungnir-Trit mode of the public implementation (`https://github.com/HKU-BAL/Gungnir`), with rates of 1.50 and 1.00 bits/nt, respectively. The low-rate configuration is obtained using the standard Gungnir mode, with a rate of 0.50 bits/nt. In all three cases, the encoded sequence length is set to 150 nts. For each configuration and bias regime, we determine the minimum coverage depth and associated read cost required for successful file retrieval at the 5% error rate. To reduce the computational burden, the reliability requirement is relaxed relative to Fig. 1: instead of requiring successful retrieval in 50 out of 50 independent trials, we require exact file recovery in a single trial at each evaluated coverage depth. The resulting coverage depth, read cost, and decoding time are reported in Fig. B below.

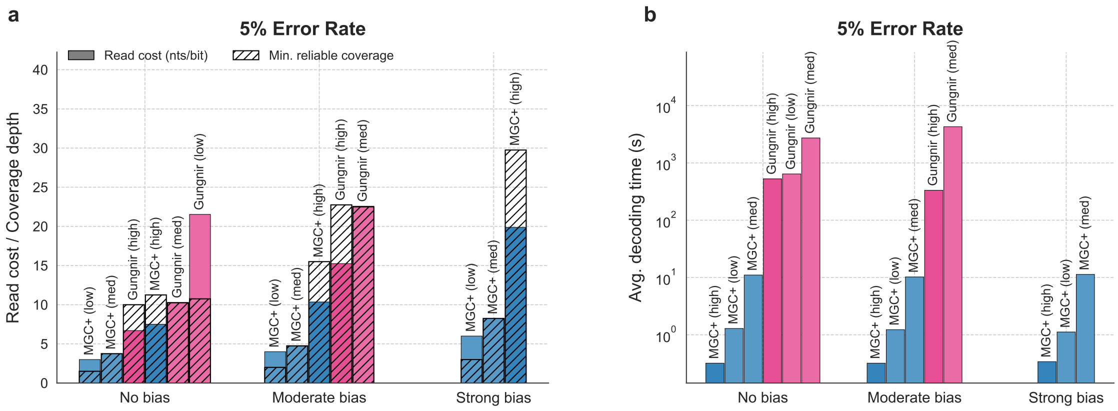


**Figure B. In silico performance comparison of DNA-MGC+ and Gungnir.** (**a**) Minimum sequencing depth and corresponding read cost required for reliable decoding at a 5% error rate under three different bias regimes. (**b**) Average decoding time measured at the minimum coverage depth required for reliable decoding.

The results show that DNA-MGC+ requires lower coverage depth and lower read cost than Gungnir across the evaluated bias regimes and code-rate classes, while also achieving orders-of-magnitude faster decoding. In the strong-bias case, Gungnir does not achieve successful retrieval within the evaluated coverage-depth range of [1, 32], which is expected since the codec is not robust to dropouts. This dropout limitation is architectural and remains regardless of the IDS error rate, while extending the comparison to higher error rates is computationally prohibitive, as previously noted for the 10% error-rate case.

#### Supplementary Note S5: Additional DNA-MGC+ results for larger file sizes and higher error rates

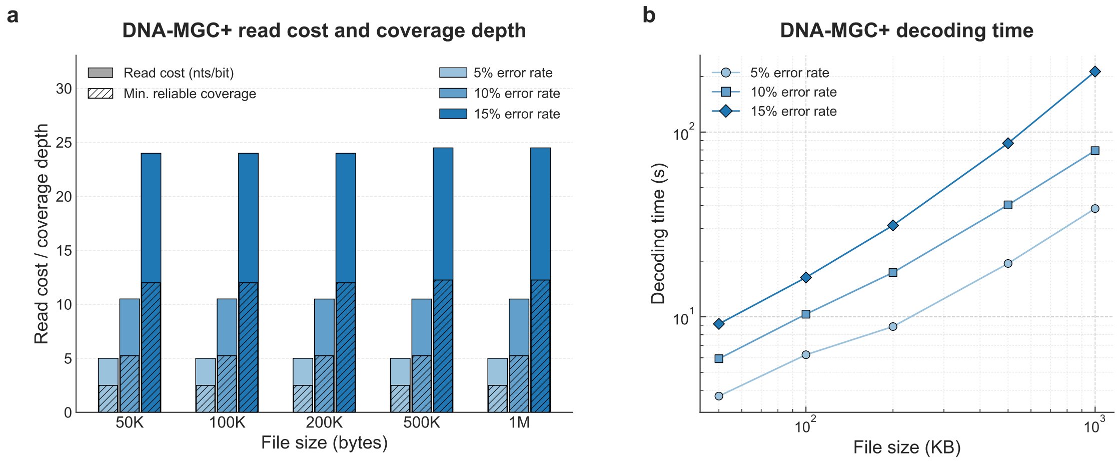


**Figure C. In silico DNA-MGC+ performance across varying file sizes and error rates under the moderate-bias scenario of the synthetic channel model.** (**a**) Minimum coverage depth required for reliable decoding and corresponding read cost (nts/bit) as a function of file size and error rate. (**b**) Average decoding time measured at the minimum coverage depth required for reliable decoding.

In Fig. 2 of the main text, we assessed the system-level scalability of DNA-MGC+ and several comparison codecs by varying the input file size from a few kilobytes to hundreds of megabytes, while fixing the error rate at 5% and considering the moderate-bias setting of the synthetic channel model. Here, we complement these results by instead fixing the file size at several representative values (50 KB, 100 KB, 200 KB, 500 KB, and 1 MB) and varying the error rate among 5%, 10%, and 15%, under the same moderate-bias setting. In Fig. C, for each file size–error rate combination, we report the minimum coverage depth required for reliable decoding together with the associated read cost, as well as the average decoding time measured at the minimum coverage depth (Fig. C b), following the same 50-out-of-50 reliability criterion and the same preprocessing pipeline (CBR clustering and Kalign alignment) used in the evaluation corresponding to Fig. 2.

The results show that, for a fixed error rate, the minimum coverage depth and read cost of DNA-MGC+ remain essentially stable as the file size increases from 50 KB to 1 MB, consistent with the trend already reported in Fig. 2 at 5% error rate. As expected, coverage depth, read cost, and decoding time all increase with the error rate; however, this increase is consistent across file sizes, indicating a favorable scalability of DNA-MGC+ over the investigated scenarios. We note that the error rates considered here, in particular 10% and 15%, are substantially higher than those typically observed in the conventional wet-lab experimental workflows and are instead intended to stress-test the scalability of the codec.

#### Supplementary Tables

**Supplementary Table S1.** Selected parameters for the DNA-MGC+ codec in the *in silico* and *in vitro* evaluations.

| Parameter | In-silico studies | | | | In-vitro experiment | | |
|---|---|---|---|---|---|---|---|
| | Low | Low (large file) | Medium | High | Opt | Design A | Design B |
| Inner Redundancy ($c_{\text{in}}$) | 12 | 10 | 6 | 0 | 6 | 6 | 6 |
| Outer code rate ($(K/(K+c_{\text{out}}))$) | 0.679 | 0.679 | 0.712 | 0.796 | 0.356 | 0.80 | 0.77 |
| Inner symbol length ($\ell_{\text{in}}$, bits) | 8 | 8 | 8 | 8 | 8 | 8 | 8 |
| Outer symbol length ($\ell_{\text{out}}$, bits) | 16 | 16 | 16 | 16 | 16 | 16 | 16 |
| Sequence index length (bits) | 16 | 16 | 16 | 16 | 16 | 16 | 16 |
| Packet index length (bits) | – | 16 | – | – | – | – | – |
| Periodic markers | Yes | Yes | No | No | No | No | Yes |
| Code rate (bits/nt) | 0.5 | 0.5 | 1 | 1.5 | 0.5 | 1.03 | 0.71 |
| Sequence length ($L_{\text{ref}}$, nts) | 152 | 152 | 148 | 152 | 148 | 124 | 122 |

**Supplementary Table S2.** Selected parameters for the DNA-Aeon codec used in the *in silico* and *in vitro* evaluations. These parameter settings are taken from Refs.[5,6].

| Parameter | In-silico studies | | | In-vitro experiment |
|---|---|---|---|---|
| | Low | Medium | High | |
| Homopolymer | 4 | 4 | 4 | 4 |
| GC-content | 0.0–1.0 | 0.0–1.0 | 0.0–1.0 | 0.0–1.0 |
| Package redundancy | 1.68 | 0.34 | 0.031 | 0.32 |
| Chunk size | 25 | 25 | 28 | 20 |
| Sync value | 4 | 4 | 8 | 4 |
| Error correction | CRC | CRC | CRC | CRC |
| Codeword length | 10 | 10 | 10 | 10 |
| CRC threshold | 3 | 3 | 3 | 3 |
| Loop | 1 | 1 | 1 | 1 |
| Finish | 0 | 0 | 0 | 0 |
| Penalty (CRC) | 0.1 | 0.1 | 0.1 | 0.1 |
| Penalty (No-Hit) | 8 | 8 | 8 | 8 |
| Code rate (bits/nt) | 0.5 | 1.01 | 1.5 | 1 |
| Sequence length (nts) | 149 | 149 | 144 | 120 |

**Supplementary Table S3.** Selected parameters for the Hedges codec used in the *in silico* and *in vitro* evaluations. The *in silico* parameter settings are taken from Refs.[5,6].

| **Parameter** | **In-silico studies** | | **In-vitro experiment** |
|---|---|---|---|
| | Low | Medium | |
| Code rate index | 3 | 1 | 3 |
| Homopolymer | 4 | 4 | 4 |
| GC window size | 12 | 12 | 12 |
| Max. GC (in window) | 8 | 8 | 8 |
| Code rate (bits/nt) | 0.65 | 1.09 | 0.61 |
| Sequence length (nts) | 148 | 147 | 126 |

**Supplementary Table S4.** Selected parameters for the RS codec used in the *in silico* evaluations. These parameter settings are taken from Refs.[5,6].

| **Parameter** | **In-silico studies** | | |
|---|---|---|---|
| | Low | Medium | High |
| Inner RS symbol length | 6 | 6 | 6 |
| Outer RS symbol length | 14 | 14 | 14 |
| Index length | 24 | 24 | 24 |
| Inner Red. symbols | 4 | 2 | 2 |
| Number of sequences | 1602 | 834 | 556 |
| Code rate (bits/nt) | 0.5 | 1 | 1.5 |
| Sequence length (nts) | 150 | 144 | 144 |

**Supplementary Table S5.** Selected parameters for the DNA-Fountain codec used in the *in silico* evaluations. These parameter settings are taken from Refs.[5,6].

| **Parameter** | **In-silico studies** | | |
|---|---|---|---|
| | Low | Medium | High |
| Alpha | 2.35 | 0.68 | 0.19 |
| Payload | 32 | 32 | 34 |
| RS length | 2 | 2 | 0 |
| Hamming distance | 100 | 100 | 100 |
| GC-content | 0.0–1.0 | 0.0–1.0 | 0.0–1.0 |
| Homopolymer | 4 | 4 | 4 |
| Delta | 0.05 | 0.1 | 0.1 |
| C-Dist | 0.1 | 0.025 | 0.025 |
| Header size | 4 | 4 | 4 |
| Code rate (bits/nt) | 0.5 | 1 | 1.5 |
| Sequence length (nts) | 152 | 152 | 152 |

**Supplementary Table S6.** Minimum reliable coverage depth for different codec configurations, error rates, and bias conditions, under the synthetic channel model (in silico). Entries are reported as unequal | equal; equal denotes $P_d = P_s = P_i$, whereas unequal denotes $P_s = 0.572P_e$, $P_d = 0.447P_e$, and $P_i = 0.026P_e$.

| Codec | Class | Code rate | Error rate | No bias | | Moderate bias | | Strong bias | |
|---|---|---|---|---|---|---|---|---|---|
| | | | | Unequal | Equal | Unequal | Equal | Unequal | Equal |
| **Aeon** | Low | 0.50 | 0.01 | 1.50 | 1.75 | 1.75 | 1.25 | 2.25 | 1.25 |
| | | | 0.05 | 3.75 | 4.50 | 4 | 5.50 | 5.25 | 8.75 |
| | Medium | 1 | 0.01 | 3 | 3.25 | 2.75 | 3.25 | 4.25 | 5 |
| | | | 0.05 | 5 | 6.25 | 5.50 | 6.25 | 10.25 | 13.25 |
| | | | 0.10 | 10 | 11.50 | 11.50 | 12 | 20.75 | 22.50 |
| | High | 1.50 | 0.01 | 6.25 | 7.25 | 9.25 | 12 | 29.50 | - |
| | | | 0.05 | 10.75 | 11.25 | 21.25 | 19.50 | - | |
| | | | 0.10 | 22.25 | - | - | | - | |
| **Fountain** | Medium | 1 | 0.01 | 12.25 | 11.25 | 23.50 | 26 | - | |
| | | | 0.05 | 25.75 | - | - | | - | |
| | High | 1.50 | 0.01 | 16 | - | - | | - | |
| **HEDGES** | Low | 0.65 | 0.01 | 3.25 | 3.50 | 4.75 | 4.75 | 11.50 | 14.25 |
| | | | 0.05 | 3.75 | 4 | 6.50 | 5.25 | 14 | 14.25 |
| | | | 0.10 | 6.25 | 6.50 | 9.25 | 9.25 | 22 | 20 |
| | | | 0.15 | 16.75 | 13 | 26.25 | 20.25 | - | |
| | Medium | 1.09 | 0.01 | 3.25 | 3.25 | 4.50 | 4.75 | 11.25 | 11 |
| | | | 0.05 | 6 | 6.25 | 9 | 9.50 | 23.25 | 23.75 |
| | | | 0.10 | 10 | 10 | 16.75 | 15.50 | - | |
| **LDPC** | Low | 0.58 | 0.01 | 2.75 | 3.75 | 3.25 | 4.50 | 5.75 | 8 |
| | | | 0.05 | 6.75 | 6.75 | 8.75 | 8.75 | 16.25 | 17.75 |
| | | | 0.10 | 18.50 | 18.50 | 24.25 | 28 | - | |
| | Low | 0.67 | 0.01 | 3.50 | 4.75 | 4.50 | 6 | 10 | 12.75 |
| | | | 0.05 | 8.75 | 8.50 | 11.50 | 12.25 | 25.50 | 27 |
| | | | 0.10 | 23 | - | 32 | - | - | |
| | Medium-Low | 0.80 | 0.01 | 7 | 8.25 | 11.50 | 14.75 | - | |
| | | | 0.05 | 15.50 | 15.25 | 27.75 | 27.25 | - | |
| **MGC+** | Low | 0.50 | 0.01 | 1.25 | 1.25 | 1.50 | 1.50 | 2.50 | 2.50 |
| | | | 0.05 | 1.75 | 1.75 | 2 | 1.75 | 3.25 | 3 |
| | | | 0.10 | 3.75 | 3.75 | 4.25 | 4.50 | 7.50 | 7.50 |
| | | | 0.15 | 12.50 | 9.75 | 16.25 | 11.50 | 30.25 | 21 |
| | Medium | 1 | 0.01 | 1.50 | 1.50 | 1.75 | 1.75 | 3.25 | 3 |
| | | | 0.05 | 4 | 4 | 5 | 4.75 | 9.25 | 8.75 |
| | | | 0.10 | 8.75 | 7 | 11.25 | 8.75 | 22.50 | 17.75 |
| | High | 1.50 | 0.01 | 4.25 | 4.50 | 5.25 | 5.50 | 11.25 | 11.25 |
| | | | 0.05 | 11.50 | 9.50 | 14.75 | 12.75 | 31.25 | 28 |
| **RS** | Low | 0.50 | 0.01 | 1 | 1.25 | 1 | 1.25 | 1.25 | 1.50 |
| | | | 0.05 | 3.25 | 3.25 | 3.25 | 3.25 | 4.25 | 4 |
| | | | 0.10 | 8.75 | 6.75 | 8.75 | 7.25 | 10.25 | 9 |
| | Medium | 1 | 0.01 | 2.75 | 3 | 3 | 3.25 | 4.50 | 5 |
| | | | 0.05 | 5.50 | 5.50 | 6.50 | 6.50 | 10.25 | 10 |
| | | | 0.10 | 13.50 | 11.50 | 16 | 14 | 25.75 | 24 |
| | High | 1.50 | 0.01 | 4.75 | 5.25 | 6.50 | 7.25 | 14.25 | 16.25 |
| | | | 0.05 | 9.25 | 8.75 | 12.50 | 12.25 | 30.50 | - |
| | | | 0.10 | 23 | 24.25 | - | | - | |
| **Stairloop** | Low | 0.49 | 0.01 | 6.25 | 6.75 | 10.25 | 16.75 | - | |
| | | | 0.05 | 12 | 15.25 | 31 | 29 | - | |
| | Medium | 0.98 | 0.01 | 4.75 | 6 | 12.25 | 10.75 | - | |
| | | | 0.05 | 12.25 | 12 | 24 | - | - | |
| | | | 0.10 | 30 | - | - | | - | |

**Supplementary Table S7.** Read cost (nts/bit) for different codec configurations, error rates, and bias conditions, under the synthetic channel model (in silico). Entries are reported as unequal | equal; equal denotes $P_d = P_s = P_i$, whereas unequal denotes $P_s = 0.572P_e$, $P_d = 0.447P_e$, and $P_i = 0.026P_e$.

| **Codec** | **Class** | **Code rate** | **Error rate** | **No bias** Unequal \| Equal | **Moderate bias** Unequal \| Equal | **Strong bias** Unequal \| Equal |
|---|---|---|---|---|---|---|
| **Aeon** | Low | 0.50 | 0.01 | 3 \| 3.50 | 3.50 \| 2.50 | 5.50 \| 2.50 |
| | | | 0.05 | 7.5 \| 9 | 8 \| 11 | 10.50 \| 17.50 |
| | Medium | 1 | 0.01 | 3 \| 3.25 | 2.75 \| 3.25 | 4.25 \| 5 |
| | | | 0.05 | 5 \| 6.25 | 5.50 \| 6.25 | 10.25 \| 13.25 |
| | | | 0.10 | 10 \| 11.50 | 11.50 \| 12 | 20.75 \| 22.51 |
| | High | 1.50 | 0.01 | 4.15 \| 4.83 | 6.15 \| 8 | 19.60 \| - |
| | | | 0.05 | 7.14 \| 7.50 | 14.12 \| 13 | - |
| | | | 0.10 | 14.78 \| - | - | - |
| **Fountain** | Medium | 1 | 0.01 | 12.23 \| 11.23 | 23.46 \| 25.95 | - |
| | | | 0.05 | 25.70 \| - | - | - |
| | High | 1.50 | 0.01 | 10.65 \| - | - | - |
| **HEDGES** | Low | 0.65 | 0.01 | 4.99 \| 5.37 | 7.29 \| 7.29 | 17.66 \| 21.88 |
| | | | 0.05 | 5.76 \| 6.14 | 9.98 \| 8.06 | 21.50 \| 21.88 |
| | | | 0.10 | 9.60 \| 9.98 | 14.20 \| 14.20 | 33.78 \| 30.71 |
| | | | 0.15 | 25.72 \| 19.96 | 40.31 \| 31.10 | - |
| | Medium | 1.09 | 0.01 | 2.97 \| 2.97 | 4.12 \| 4.35 | 10.30 \| 10.07 |
| | | | 0.05 | 5.49 \| 5.72 | 8.24 \| 8.69 | 21.28 \| 21.74 |
| | | | 0.10 | 9.15 \| 9.15 | 15.33 \| 14.18 | - |
| **LDPC** | Low | 0.58 | 0.01 | 4.71 \| 6.42 | 5.57 \| 7.71 | 9.85 \| 13.70 |
| | | | 0.05 | 11.56 \| 11.56 | 14.99 \| 14.99 | 27.83 \| 30.40 |
| | | | 0.10 | 31.68 \| 31.68 | 41.53 \| 47.95 | - |
| | Low | 0.67 | 0.01 | 5.20 \| 7.05 | 6.68 \| 8.91 | 14.84 \| 18.93 |
| | | | 0.05 | 12.99 \| 12.62 | 17.07 \| 18.18 | 37.85 \| 40.08 |
| | | | 0.10 | 34.14 \| - | 47.50 \| - | - |
| | Medium-Low | 0.80 | 0.01 | 8.79 \| 10.36 | 14.45 \| 18.53 | - |
| | | | 0.05 | 19.47 \| 19.16 | 34.86 \| 34.23 | - |
| **MGC+** | Low | 0.50 | 0.01 | 2.50 \| 2.50 | 3 \| 3 | 5 \| 5 |
| | | | 0.05 | 3.50 \| 3.50 | 4 \| 3.50 | 6.50 \| 6 |
| | | | 0.10 | 7.51 \| 7.51 | 8.51 \| 9.01 | 15.01 \| 15.01 |
| | | | 0.15 | 25.02 \| 19.51 | 32.52 \| 23.02 | 60.54 \| 42.03 |
| | Medium | 1 | 0.01 | 1.50 \| 1.50 | 1.75 \| 1.75 | 3.25 \| 3 |
| | | | 0.05 | 4 \| 4 | 5 \| 4.75 | 9.26 \| 8.76 |
| | | | 0.10 | 8.76 \| 7.01 | 11.26 \| 8.76 | 22.50 \| 17.77 |
| | High | 1.50 | 0.01 | 2.83 \| 3 | 3.50 \| 3.67 | 7.50 \| 7.50 |
| | | | 0.05 | 7.67 \| 6.33 | 9.83 \| 8.50 | 20.84 \| 18.67 |
| **RS** | Low | 0.50 | 0.01 | 2 \| 2.50 | 2 \| 2.50 | 2.50 \| 3 |
| | | | 0.05 | 6.51 \| 6.51 | 6.51 \| 6.51 | 8.51 \| 8.01 |
| | | | 0.10 | 17.51 \| 13.51 | 17.52 \| 14.51 | 20.52 \| 18.02 |
| | Medium | 1 | 0.01 | 2.76 \| 3.01 | 3.01 \| 3.26 | 4.51 \| 5.01 |
| | | | 0.05 | 5.51 \| 5.51 | 6.51 \| 6.51 | 10.27 \| 10.02 |
| | | | 0.10 | 13.53 \| 11.52 | 16.03 \| 14.03 | 25.80 \| 24.05 |
| | High | 1.50 | 0.01 | 3.17 \| 3.51 | 4.34 \| 4.84 | 9.52 \| 10.85 |
| | | | 0.05 | 6.18 \| 5.84 | 8.35 \| 8.18 | 20.37 \| - |
| | | | 0.10 | 15.36 \| 16.20 | - | - |
| **Stairloop** | Low | 0.49 | 0.01 | 12.71 \| 13.73 | 20.85 \| 34.06 | - |
| | | | 0.05 | 24.40 \| 31.01 | 63 \| 58.98 | - |
| | Medium | 0.98 | 0.01 | 4.83 \| 6.10 | 12.46 \| 10.93 | - |
| | | | 0.05 | 12.46 \| 12.20 | 24.40 \| - | - |
| | | | 0.10 | 30.51 \| - | - | - |

**Supplementary Table S8.** Average decoding time (seconds), measured at minimum reliable coverage depth, for different codec configurations, error rates, and bias conditions, under the synthetic channel model (in silico). Entries are reported as unequal | equal; equal denotes $P_d = P_s = P_i$, whereas unequal denotes $P_s = 0.572P_e$, $P_d = 0.447P_e$, and $P_i = 0.026P_e$.

| Codec | Class | Code rate | Error rate | No bias<br>Unequal \| Equal | Moderate bias<br>Unequal \| Equal | Strong bias<br>Unequal \| Equal |
|---|---|---|---|---|---|---|
| Aeon | Low | 0.50 | 0.01 | 1715.46 \| 4141.76 | 1721.73 \| 3571.20 | 925.15 \| 2626.28 |
| | | | 0.05 | 5931.75 \| 6544.29 | 6539.69 \| 6784.33 | 6408.26 \| 6556.18 |
| | Medium | 1 | 0.01 | 496.54 \| 1314.22 | 587.18 \| 1395.69 | 451.27 \| 1119.79 |
| | | | 0.05 | 1985.66 \| 1285.73 | 2802.16 \| 2727.98 | 2503 \| 2130.38 |
| | | | 0.10 | 3035.07 \| 3452.06 | 4020.46 \| 4669.22 | 4613.66 \| 5347.59 |
| | High | 1.50 | 0.01 | 66.19 \| 80.08 | 74.64 \| 75.92 | 61.05 \| - |
| | | | 0.05 | 58.88 \| 71.48 | 62.67 \| 91.97 | - |
| | | | 0.10 | 1494.41 \| - | - | - |
| Fountain | Medium | 1 | 0.01 | 0.61 \| 0.62 | 0.59 \| 0.61 | - |
| | | | 0.05 | 0.62 \| - | - | - |
| HEDGES | Low | 0.65 | 0.01 | 10.02 \| 6.68 | 7.56 \| 6.68 | 6.71 \| 6.56 |
| | | | 0.05 | 11.33 \| - | 8.73 \| 10.80 | 7.80 \| 8.10 |
| | | | 0.10 | 52.70 \| 52 | 55.27 \| 58.26 | 64.61 \| 72.21 |
| | | | 0.15 | 2248.37 \| 1510.76 | 2921.46 \| 1906.64 | - |
| | Medium | 1.09 | 0.01 | 5.18 \| 5.49 | 5.34 \| 5.90 | 4.66 \| 5.06 |
| | | | 0.05 | 13.20 \| 13 | 13.33 \| 12.83 | 10.60 \| 12.75 |
| | | | 0.10 | 35.01 \| 42.23 | 40.12 \| 47.39 | - |
| LDPC | Low | 0.58 | 0.01 | 164.53 \| 151.25 | 161.02 \| 174.68 | 155.83 \| - |
| | | | 0.05 | 171.66 \| 183.83 | 168.34 \| 177.13 | 163.43 \| 169.40 |
| | | | 0.10 | 176.77 \| 192.31 | 174.87 \| 192.49 | - |
| | Low | 0.67 | 0.01 | 138.75 \| 157.88 | 147.18 \| 145.56 | 141.58 \| 134.17 |
| | | | 0.05 | 147.44 \| 157 | 112.61 \| 154.67 | 138.27 \| 122.01 |
| | | | 0.10 | 101 \| - | 129.22 \| - | - |
| | Medium-Low | 0.80 | 0.01 | 123.31 \| 135.56 | 131.28 \| 132.21 | - |
| | | | 0.05 | 125.03 \| 130.78 | 124.84 \| 125.98 | - |
| MGC+ | Low | 0.50 | 0.01 | 38.51 \| 44.41 | 37.97 \| 42.63 | 39.08 \| 43.46 |
| | | | 0.05 | 62.64 \| 85.75 | 58.18 \| 80.59 | 54.74 \| 70.32 |
| | | | 0.10 | 218.19 \| 221.23 | 216.29 \| 224.36 | 193.94 \| 216.05 |
| | | | 0.15 | 3624.93 \| 2466.74 | 4142.30 \| 2644.65 | 5687.49 \| 3541.33 |
| | Medium | 1 | 0.01 | 40.88 \| 139.09 | 37.56 \| 116.78 | 31.34 \| 90.04 |
| | | | 0.05 | 594.15 \| 1079.25 | 529.17 \| 1063.09 | 657.98 \| 886.38 |
| | | | 0.10 | 1223.52 \| 1774.63 | 1046 \| 1764.74 | 1599.47 \| 1764.34 |
| | High | 1.50 | 0.01 | 6.65 \| 6.78 | 6.88 \| 6.36 | 6.77 \| 9.79 |
| | | | 0.05 | 6.57 \| 6.49 | 6.67 \| 6.46 | 6.61 \| 6.22 |
| RS | Low | 0.50 | 0.01 | 2210.36 \| 2625.96 | 2410.36 \| 2652.63 | 2347.29 \| 2603.80 |
| | | | 0.05 | 2251.17 \| 2466.59 | 2419.03 \| 2601.99 | 2327.53 \| - |
| | | | 0.10 | 2224.46 \| 2566.19 | 2496.38 \| 2469.72 | 2644.44 \| 2507.66 |
| | Medium | 1 | 0.01 | 169.56 \| 180.53 | 189.42 \| 206.43 | 197 \| 200.26 |
| | | | 0.05 | 191.69 \| 191.59 | 191.16 \| 190.71 | 198.28 \| 207.99 |
| | | | 0.10 | 203.49 \| 209.68 | - \| 211.17 | 205.81 \| 211.01 |
| | High | 1.50 | 0.01 | 8.20 \| 6.47 | 8.16 \| 6.95 | 8.48 \| 8.21 |
| | | | 0.05 | 8.64 \| 9.19 | 9.44 \| 9.67 | 7.86 \| - |
| | | | 0.10 | 8.64 \| 8.43 | - | - |
| Stairloop | Low | 0.49 | 0.01 | 606.85 \| 689.71 | 1023.71 \| 1688.34 | - |
| | | | 0.05 | 1632.23 \| 1828.34 | 4598.35 \| 3464.44 | - |
| | Medium | 0.98 | 0.01 | 247.24 \| 313.80 | 590.52 \| 552.66 | - |
| | | | 0.05 | 882.21 \| 731.87 | 1628.82 \| - | - |
| | | | 0.10 | 2747.86 \| - | - | - |

**Supplementary Table S9.** Performance metrics for different codec configurations under the low-fidelity experimentally derived error and bias profiles (in silico) for a fixed physical redundancy of 100×.

| Codec | Class | Code rate | Sequencing depth | Read cost (nts/bit) | Avg. decoding time (s) |
|---|---|---|---|---|---|
| **Aeon** | Low | 0.50 | 12.25 | 24.49 | 2440.82 |
| | Medium | 1 | 9.75 | 9.75 | 1533.13 |
| **HEDGES** | Low | 0.65 | 16.75 | 25.72 | 7.25 |
| | Medium | 1.09 | 20.25 | 18.53 | 10.65 |
| **MGC+** | Low | 0.50 | 2.50 | 5 | 41.82 |
| | Medium | 1 | 4.50 | 4.50 | 666.87 |
| | Optimized | 0.5 | 1 | 2 | 1251.82 |
| **RS** | Low | 0.50 | 3.25 | 6.51 | 2352.07 |
| | Medium | 1 | 12.75 | 12.77 | 210.18 |

**Supplementary Table S10.** Performance metrics for different codec configurations under the low-fidelity experimentally derived error and bias profiles (in silico) for a fixed sequencing depth of 30×.

| Codec | Class | Code rate | Physical redundancy | Write cost (nts/bit) | Avg. decoding time (s) |
|---|---|---|---|---|---|
| **Aeon** | Low | 0.50 | 13.50 | 26.99 | 3223.77 |
| | Medium | 1 | 14.50 | 14.51 | 1584.41 |
| **HEDGES** | Low | 0.65 | 15.75 | 24.19 | 8.73 |
| | Medium | 1.09 | 30.75 | 28.14 | 8.12 |
| **MGC+** | Low | 0.50 | 2.25 | 4.50 | 40.78 |
| | Medium | 1 | 4.25 | 4.25 | 642.49 |
| | Optimized | 0.50 | 1 | 2 | 1190.30 |
| **RS** | Low | 0.50 | 5 | 10.01 | 2492.93 |
| | Medium | 1 | 20.75 | 20.79 | 214.40 |

**Supplementary Table S11.** Minimum reliable sequencing depth, read cost, and average decoding time (serial and parallel), under Illumina sequencing and Nanopore sequencing with different basecallers (in vitro).

| Sequencing method | Codec | Min. reliable sequencing depth | Code rate (bits/nt) | Read cost (nts/bit) | Avg. decoding time (s) 1 core \| 8 cores |
|---|---|---|---|---|---|
| Illumina | MGC+ (A, unfiltered) | 2.75 | 1.032 | 2.66 | 33.34 \| 8.08 |
| | MGC+ (A, filtered) | 2.50 | 1.032 | 2.42 | 207.74 \| 13.98 |
| | MGC+ (B, unfiltered) | 2.75 | 0.706 | 3.90 | 58.66 \| 7.51 |
| | MGC+ (B, filtered) | 2.25 | 0.706 | 3.19 | 438.83 \| 20.09 |
| | HEDGES | 9.50 | 0.610 | 15.57 | 32.48 \| – |
| | Aeon | 3.00 | 1.000 | 3.00 | 571.12 \| 119.23 |
| Nanopore (dorado-sup) | MGC+ (A, unfiltered) | 4.00 | 1.032 | 3.88 | 114.65 \| 14.53 |
| | MGC+ (A, filtered) | 3.50 | 1.032 | 3.39 | 193.58 \| 22.43 |
| | MGC+ (B, unfiltered) | 3.00 | 0.706 | 4.25 | 63.60 \| 7.98 |
| | MGC+ (B, filtered) | 2.75 | 0.706 | 3.90 | 406.03 \| 20.89 |
| | HEDGES | 7.50 | 0.610 | 12.30 | 31.99 \| – |
| | Aeon | 5.00 | 1.000 | 5.00 | 2494.72 \| 305.97 |
| Nanopore (dorado-hac) | MGC+ (A, unfiltered) | 5.25 | 1.032 | 5.09 | 154.66 \| 19.07 |
| | MGC+ (A, filtered) | 4.25 | 1.032 | 4.12 | 310.40 \| 27.45 |
| | MGC+ (B, unfiltered) | 3.50 | 0.706 | 4.96 | 67.05 \| 8.22 |
| | MGC+ (B, filtered) | 3.25 | 0.706 | 4.60 | 471.67 \| 21.98 |
| | HEDGES | 8.25 | 0.610 | 13.52 | 36.28 \| – |
| | Aeon | 6.50 | 1.000 | 6.50 | 3256.71 \| 355.40 |
| Nanopore (dorado-fast) | MGC+ (A, unfiltered) | 10.75 | 1.032 | 10.42 | 389.59 \| 40.42 |
| | MGC+ (A, filtered) | 9.25 | 1.032 | 8.96 | 502.40 \| 44.36 |
| | MGC+ (B, unfiltered) | 6.75 | 0.706 | 9.56 | 86.09 \| 9.76 |
| | MGC+ (B, filtered) | 6.50 | 0.706 | 9.21 | 503.16 \| 23.76 |
| | HEDGES | 11.50 | 0.610 | 18.85 | 85.37 \| – |
| | Aeon | 13.00 | 1.000 | 13.00 | 5071.89 \| 537.22 |

### Supplementary Figures

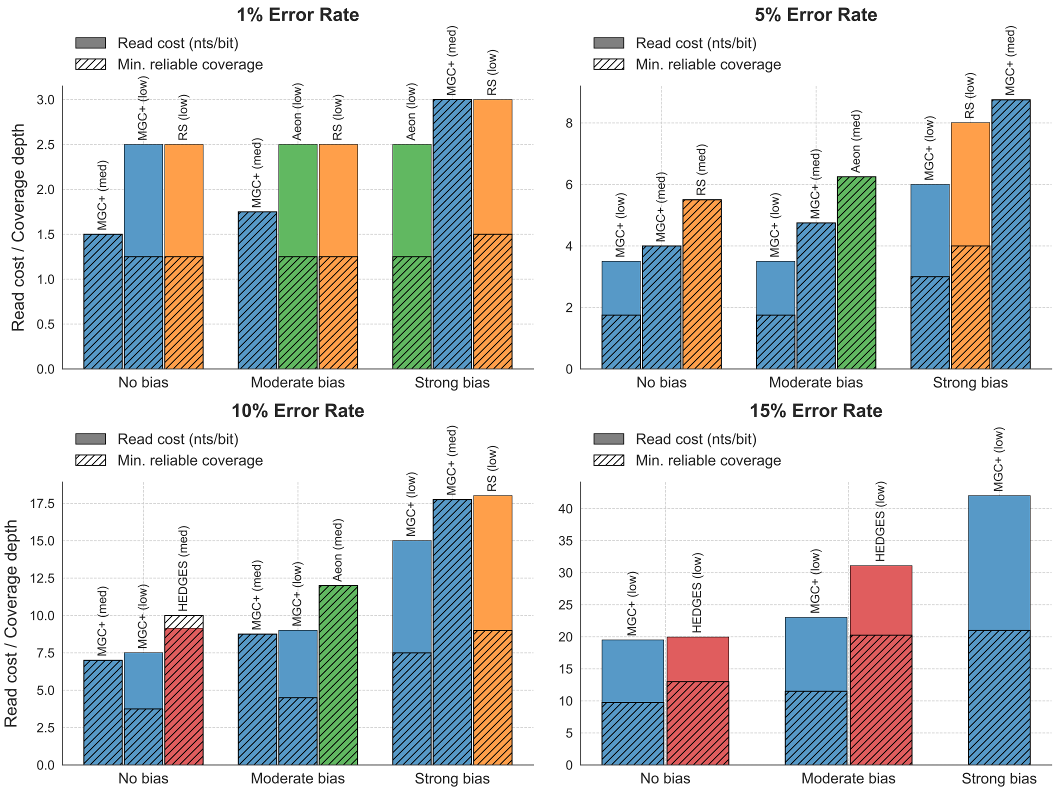


**Supplementary Figure S1.** Minimum coverage depth and associated read cost achieved by the three best-performing codecs across the different bias and error combinations for the case where $P_s = P_d = P_i$.

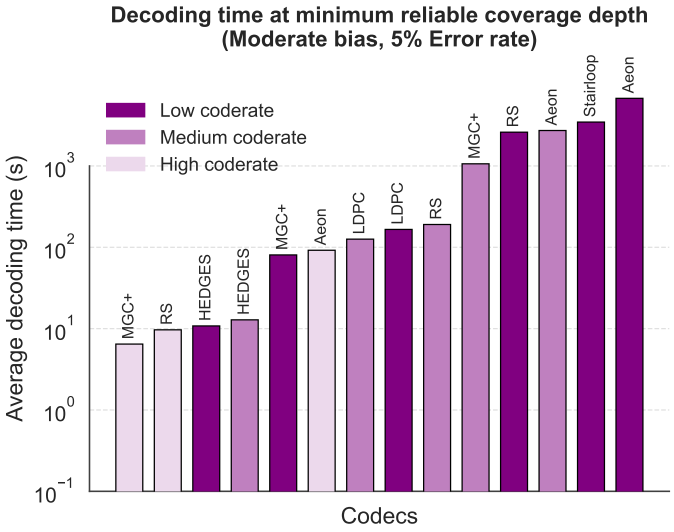


**Supplementary Figure S2.** Average decoding times for the moderate bias and 5% error rate scenario, measured at the minimum coverage depth required for reliable decoding for the case where $P_s = P_d = P_i$.

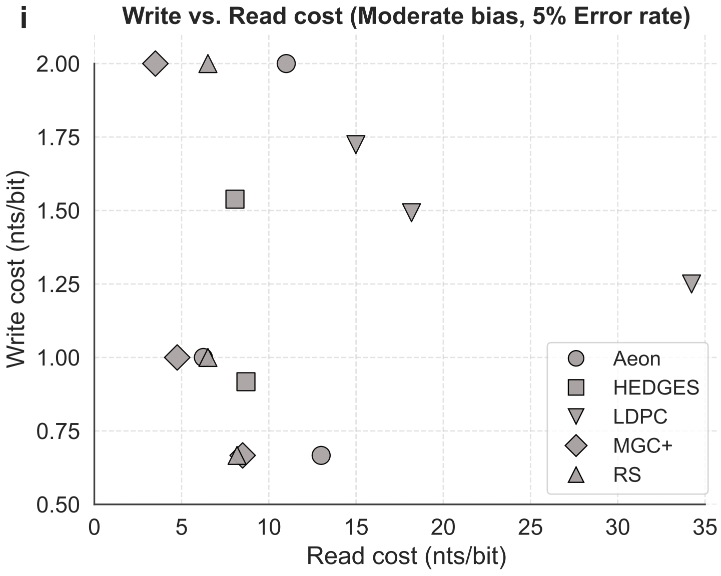


**Supplementary Figure S3.** Trade-off between write cost and read cost for the moderate bias and 5% error rate scenario for the case where $P_s = P_d = P_i$.

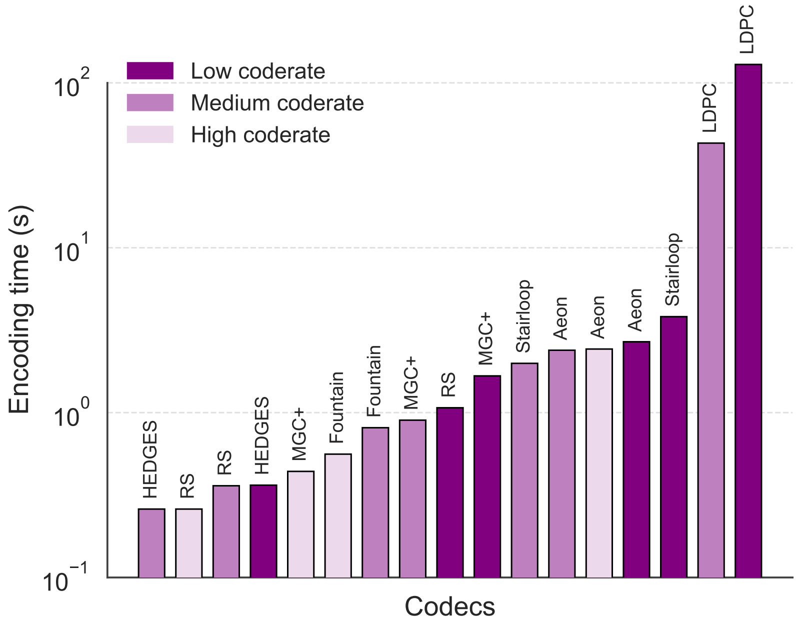


**Supplementary Figure S4.** Encoding times required by different codecs for a 15KB file.

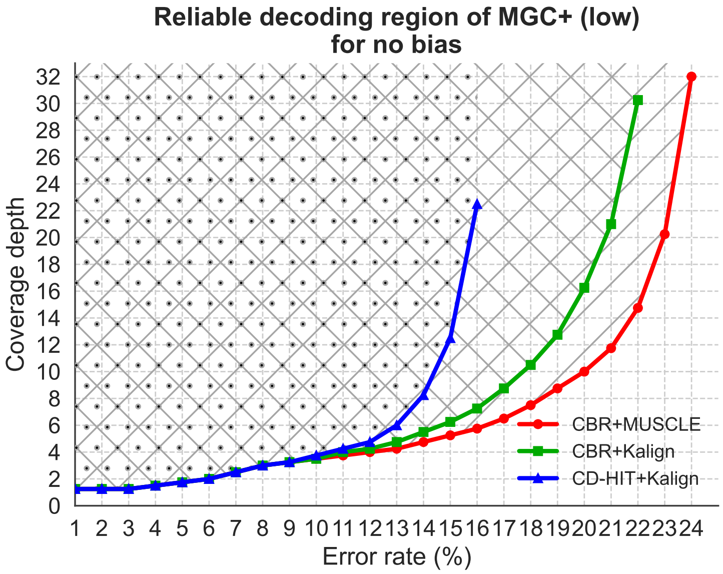


**Supplementary Figure S5.** Reliable decoding region of DNA-MGC+ (low-rate) for the no-bias case.

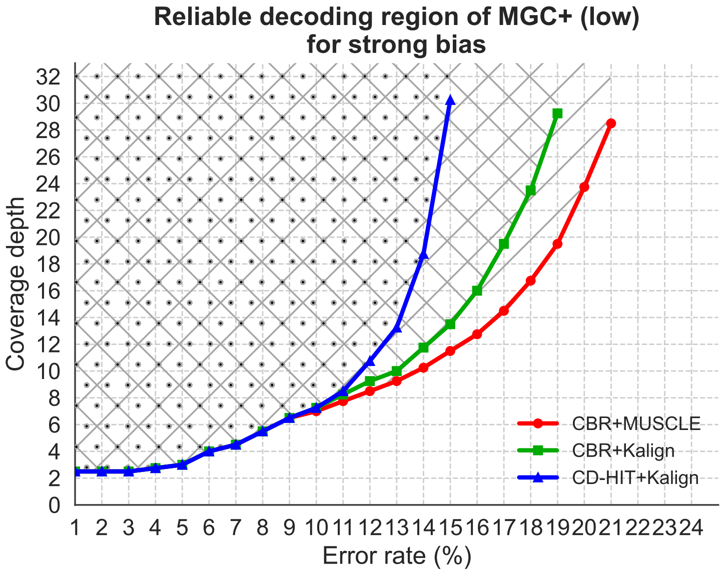


**Supplementary Figure S6.** Reliable decoding region of DNA-MGC+ (low-rate) for the strong-bias case.

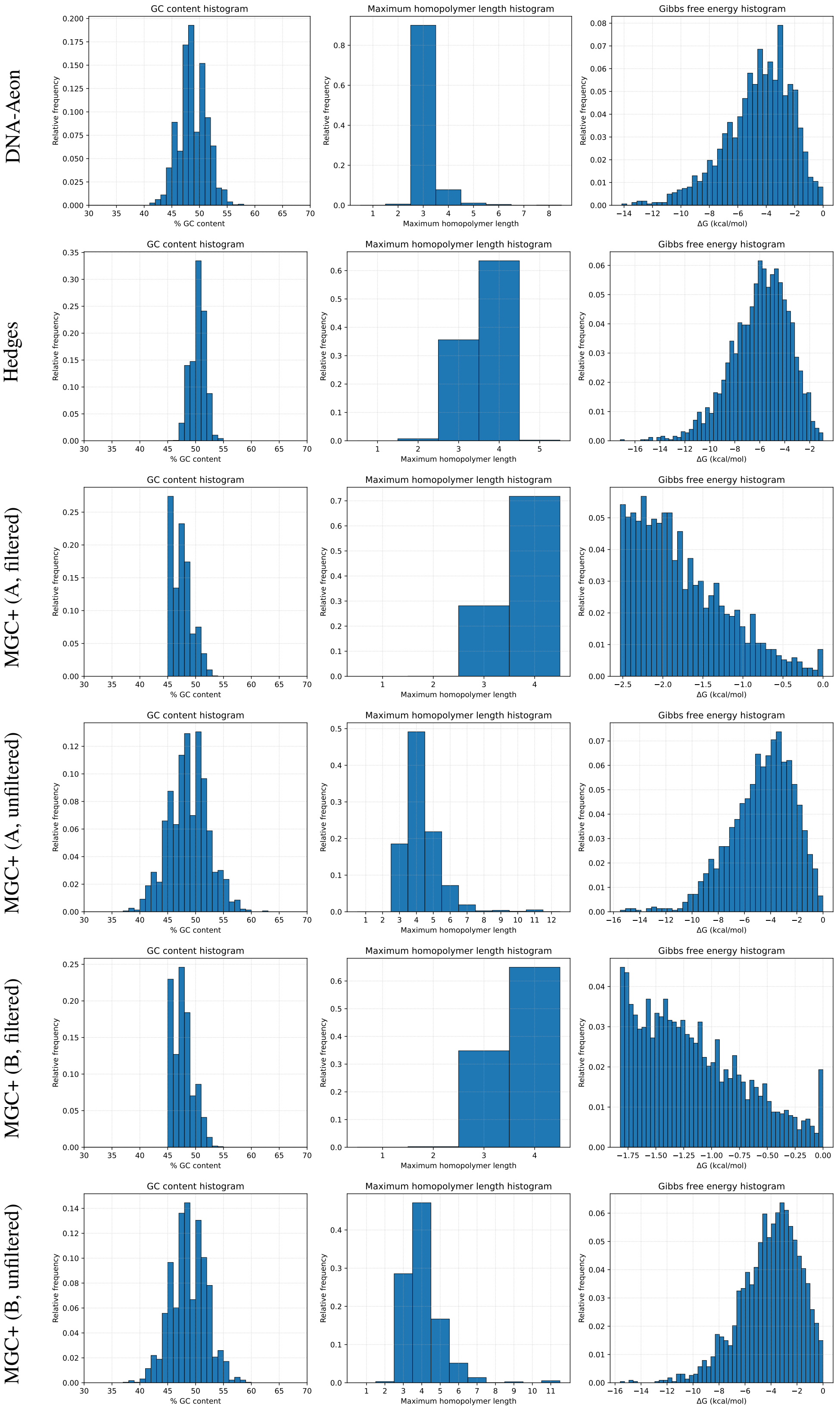


**Supplementary Figure S7.** Normalized histograms of GC content, maximum homopolymer length, and Gibbs free energy ($\Delta G$) for the encoded sequences of the different configurations included in the *in vitro* experiment.

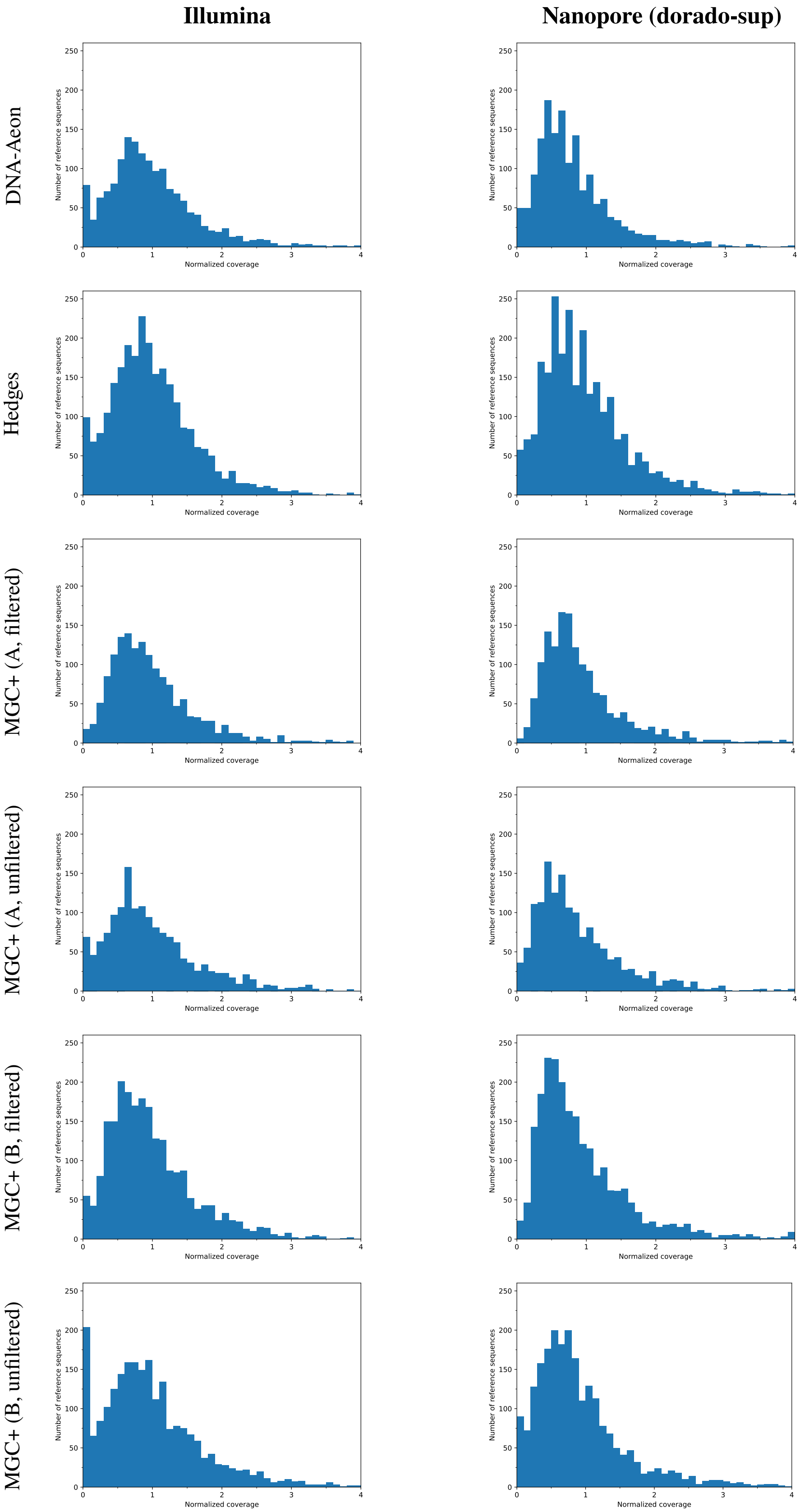


**Supplementary Figure S8.** Coverage distributions derived from the Illumina and Nanopore (dorado-sup) sequencing data in the *in vitro* experiment, normalized by the mean coverage of each setting.
